# Assessing planktivorous fish as vectors of a plankton parasite

**DOI:** 10.64898/2026.07.09.737450

**Authors:** Nikolaos-Dimitrios Lampadaridis, Carlos Manuel Herrera-Castillo, Dieter Ebert

**Affiliations:** Department of Environmental Sciences, Zoology, University of Basel, Vesalgasse 1, 4051 Basel, Switzerland

## Abstract

According to the "healthy herd hypothesis," which posits that predators can reduce parasite prevalence in prey populations by preferentially removing infected individuals, predators are often thought to regulate disease in their prey. However, predators may also act as disease vectors, facilitating the spread of parasites. Here we investigate whether stickleback fish (*Gasterosteus aculeatus*) act as vectors in the transmission of the obligate bacterial parasite *Pasteuria ramosa* to its *Daphnia* host, a widespread freshwater zooplanktor. We fed infected *D. magna* to sticklebacks and subsequently analysed faecal samples for the presence, viability, and infectivity of parasite transmission stages (= spores). In the approximately 60% of consumed spores that we recovered from fish faeces, we found no reduction in infectivity to *D. magna*. Nor did spores associated with sloppy feeding show reduced infection rates. Thus, by consuming infected hosts, the fish does not eliminate the parasite, but rather may contribute to the spread and persistence of *P. ramosa* in natural populations, potentially influencing parasite dynamics in natural freshwater ecosystems.

## Introduction

Parasites play pivotal roles in shaping the structure, dynamics, and function of ecosystems (Lafferty et al., 2006; Brian et al., 2022). In aquatic environments, parasite influence is integral yet often overlooked (Selbach et al., 2020), as their interactions with hosts can regulate species abundance, affect food web dynamics, and drive evolutionary processes (Johnson et al., 2010; Brunner et al., 2017; Timi & Poulin, 2020). Some aquatic parasites act as ecosystem engineers, altering host behaviour or morphology in ways that influence habitat structure or nutrient cycling (Wood et al., 2007; Mischler et al., 2016). Infected hosts often become more easily preyed upon, as the may be more visible or have an impaired escape responses, and thereby altering predator–prey dynamics and potentially disrupting trophic interactions. When infected with a bacterial parasite, zooplankton, for instance, become more visible to predatory fish such as bluegill sunfish, resulting in higher predation rates (Wale et al. 2021).

Given these effects of predators, it is important to understand how parasites are transmitted and maintained in food webs, and the role that host predation plays in influencing host–parasite interactions. Because predators have been shown to target vulnerable individuals, those that are sick or slow (Hudson et al., 1992; Genovart et al., 2010), they potentially interrupt parasite transmission and evolution by removing infectious hosts from the population (Packer et al., 2003; Duffy et al., 2005; Gallagher et al., 2019), which is in the centre of the “healthy herd hypothesis”. This hypothesis states that predators regulate disease by targeting infected individuals (Packer et al., 2003; Richards et al., 2023). By acting as terminal sinks, predators are thought to reduce parasite prevalence and outbreak risk (Duffy et al., 2005; Gutierrez et al., 2022). A core assumption for this hypothesis is that parasites perish when infected hosts are consumed. However, this assumption may not always hold true, as parasites may survive the passage through the predator’s gut and then spread from the predator’s faeces.

Indeed, parasites may have thick-walled, well protected transmission stages, which may facilitate survival during a gut passage, remain infectious, and be redeposited into the environment, in which case predators could facilitate parasite persistence and spread rather than reducing it (Duffy, 2009). Such adaptations may play an active role in facilitating parasite dissemination by serving as mechanical or trophic vectors (Duffy et al., 2005; Hall et al., 2007; Marcogliese, 2022). Here we focus on predator-facilitated parasite spread by investigating whether a common planktivorous fish can function as a vector for an obligate bacterial parasite within an aquatic zooplankton system. In a series of experiments, we explore the potential of transmission stage (=spores) excretion with the faeces and other feeding-associated transmission routes, such as the waterborne transmission of spores released during the sloppy feeding of the predator, i.e., the unintentional release of prey tissue and parasite spores during prey handling of the predator.

Our host in this study is the freshwater zooplankton *Daphnia magna* (Crustacea: Cladocera) (Ebert et al., 2016; Cornetti et al., 2019), a species that occupies a wide range of fresh- and brackish-water habitats, including lakes, ponds and coastal rock pools (Ebert, 2022). Often a dominant member of the zooplankton community, it plays a vital role in freshwater food webs. *D. magna* filter-feeds on suspended particles of mainly single-celled algae, rendering it susceptible to ingesting parasite spores such as those of the bacterial pathogen *Pasteuria ramosa* (Ebert, 2022, 2025), our parasite model in this study. The spores of this obligate endoparasite are transmitted exclusively horizontally and characterized by a thick outer layer called an exosporium (Ebert et al., 1996). Upon contact with the host, the parasite sheds its exosporium and enters the host’s body cavity where it proliferates and produces spores, which are ultimately resuspended into the water column after the host’s death (Ebert et al., 2004). Infection severely impacts host fitness, inducing reproductive castration, gigantism, and premature mortality (Ebert et al., 2004). A well visible symptom of infection is a change in host coloration, with infected individuals losing transparency and becoming opaque reddish-brown (Ebert, 2005), potentially increasing their detectability by visually hunting predators. Indeed, Wale et al. (2021) found that planktivorous fish preferentially target infected *Daphnia*, likely due to increased detectability.

In many lakes, *Daphnia* is a major dietary component for planktivorous fish (Galbraith, 1967; Duffy et al., 2005; Ebert, 2005; Duffy, 2007; Lampert and Sommer, 2007). One such predator is the three-spined stickleback (*Gasterosteus aculeatus*), the predator system in this study. This small teleost fish has a broad coastal and freshwater distribution across the Northern Hemisphere (Reimchen, 1994; Woottoon, 2009; Reid et al., 2021). Sticklebacks are well-documented consumers of *Daphnia* and exhibit varied diets and high phenotypic plasticity (Bretzel et al., 2021), making them a suitable model for investigating predator–prey–parasite interactions.

Here we examine how interactions between *D. magna*, the bacterial parasite *P. ramosa* and their natural three-spined stickleback predator may influence parasite transmission. Our research objectives were: (1) to determine whether *P. ramosa* spores can pass through the gut of the fish predator, (2) to assess whether spores remain viable and infective after excretion from the gut and (3) to test the infectivity of spores released into the water either via predator excretion or feeding activity (sloppy feeding). We quantified the number of transmission stages at each step of the predation process and compared it to the infection success of spores transmitted directly from infected *Daphnia* to clarify the ecological role of fish-mediated parasite transmission.

## Material and Methods

### Experimental design and culture conditions

For this experiment, we used the *Daphnia magna* host clone HU-HO-2 (origin: Bogarzótó, Hungary, GPS: 46.8, 19.133) and the *P. ramosa* parasite clone C1 (Moscow, Russia, GPS: 55.763, 37.582) (Bento et al. 2017). Host cultures were maintained in ADaM (Artificial Daphnia Medium, Klüttgen et al., 1994, as modified by Ebert, 1998) under standardized conditions (20 °C, 16:8 h light:dark photoperiod), following protocols described in Hall & Ebert (2012) and Ebert et al. (1993). Adult females were kept in 350-mL jars and fed daily with unicellular *Tetradesmus* sp. algae. Culture density, transfer frequency, and food levels were adjusted across reproductive stages to ensure high reproduction and consistent host quality. Offspring from the first clutch were not included, as they are much smaller than later offspring.

The three-spined sticklebacks used in this study were sourced from an established laboratory culture at the University of Basel, which had been founded from fish collected at a coastal lagoon breeding site at Loch an Duin (57°38′32.96″ N, 7°12′31.76″ W) (Herrera-Castillo et al., 2025). In total, 20 fish (both males and females) were used. Five fish served as negative controls, five fish were assigned to the experimental replicates, and ten were included as companion fish to comply with animal welfare regulations (social housing to reduce stress). Each replicate fish was placed together with a companion fish in a tank that contained 2.4 L of tap water. Air stones ensured oxygenation and water circulation. Tanks were maintained at 14 °C under a 16:8 h light:dark photoperiod. Prior to the start of the infection experiment, fish were fed daily with frozen bloodworms or mosquito larvae.

### Gut passage pilot experiment

To determine an appropriate collection schedule for faecal samples in the main experiment, we conducted a pilot study to estimate gut passage time in the three-spined stickleback. One male and one female fish were randomly selected and placed in an experimental tank. Both fish were starved for 24 hours, after which they were each fed an equal amount of mosquito larvae. A camera (GoPro Hero 4) was used in time-lapse mode (10-second intervals) to record faecal deposition over three days. Faecal material was collected daily, and the video material was analysed to determine the timing of droppings for each fish. Across the three-day period, both fish produced an equal number of droppings (n = 7 each, 14 total), with four faecal events on day 1, nine on day 2 and one on day 3 (Supplementary Fig. 1). Based on these results, we collected faeces from exposed fish in 2-hour intervals over a 48-hour period in the main experiment.

### Production of infected *Daphnia*

Juvenile *D. magna* individuals age 3 to 5 days were exposed to the parasite (*P. ramosa* C) following the infection protocol described by Routtu and Ebert (2015). The exposure was carried out in 100-mL jars, filled with 20 mL of ADaM, each containing five juvenile *Daphnia*. To each jar we added a concentration of approximately 80,000 parasite spores per host individual (400000 spores per jar). After one week the jars were topped up with ADaM. In these conditions, the majority of *Daphnia* became infected and produced large quantities of parasite transmission stages. Throughout this infection phase, each *D. magna* was fed 10 million green algae cells daily and transferred to fresh medium once or twice per week. Infections became visible within 2-3 weeks after exposure. Infected individuals were maintained under controlled conditions, so parasites could produce large numbers of fully mature spores in the host’s body cavity. These mature, infected *Daphnia* were subsequently used as prey in the predation experiments.

### Predation experiment and spore recovery after gut passage

Prior to the predation experiment, we estimated the total number of spores in an infected *Daphnia*. We selected hosts of similar sizes and stages, each from a different culture jar to avoid potential jar-effects. Five times, five *Daphnia* were pooled in a 1.5-mL Eppendorf tube, homogenized in 1 mL water, and the spore concentration was quantified with a hemocytometer (Neubauer improved counting chamber, depth: 0.1 mm; square size: 0.05 mm). The average spore count per pool of five infected *Daphnia*, across five replicate pools, was 13.0 ± 0.45 million spores (mean ± SE).

For the main spore recovery experiment, we used five experimental tanks, each containing a pair of sticklebacks (one replicate and one companion fish) that had been starved for 24 hours to reduce prior gut content. One fish from each pair was temporarily isolated and offered five infected *Daphnia*, while its companion received bloodworms. This feeding regime allowed us to attribute any spores recovered in faeces specifically to the *Daphnia* exposed individual. Faecal pellets were collected with a glass pipette at 2-hour intervals over a 48-hour period to minimize potential loss of spores due to the breakdown or dispersal of the faecal material. They were then transferred into 1.5-mL Eppendorf tubes, and homogenized in 1 mL of water. Spores were quantified with a hemocytometer.

### Infection experiment with gut passaged spores

#### Feeding regime for infection experiment

We conducted a second predation experiment, this time offering five infected *D. magna* to both of the two fish and closely monitoring their consumption. After ingesting all prey, the fish remained in the feeding tanks for an additional 30 minutes to allow for the potential expulsion of ingested material (sloppy feeding). We then placed both the replicate and the companion fish together in another experimental tank for the main phase of the experiment and stored the water from the fish’s *Daphnia* feeding tank in a cold room at 8 °C for later analysis of sloppy feeding effects. For two consecutive days post feeding, we collected faeces from the fed fish by transferring them to clean tanks and retrieving faecal pellets from the bottom of their tanks using a glass pipette. We placed the faecal samples in 1.5-mL Eppendorf tubes with excess water carefully removed and stored them at 4 °C. Once collection was completed, we returned the fish to their original tanks, maintaining the existing water to allow spores to accumulate over time. After the collection period, we also collected and stored this water under the same conditions for analysis. We replicated this procedure for the control group, which was fed uninfected *Daphnia*, to confirm that faeces do not contain spores in the absence of infected prey. This experiment resulted in three potential sources of spores: spores released during feeding (sloppy feeding), spores contained in the faecal matter, and other (non faecal) spores released into the water by the fish post feeding. We quantified spores in the faecal sample with the hemocytometer and subsequently used them in infection assays to determine spore infectivity. Spores released into the water were not quantified, but the water was tested for the presence of infectious spores (see below).

#### Infection success of gut passaged spores

The spores collected from stickleback faeces were quantified as above and used to initiate an infection experiment with three exposure doses per animal: low (5,000 spores), medium (30,000 spores), and high (180,000 spores). This wide range was chosen, as we had no prior knowledge which dose would yield an appropriate result. In total, we used 380 juvenile *D. magna* in this experiment, produced as outlined above. We collected and pooled cohorts of juveniles born within a 24-hour period, into a 1.5-L jar and the following day distributed these 2-day-old juveniles individually into 380 100-mL jars, each filled with 20 mL of ADaM.

The experiment consisted of two infection treatments. For the control treatment, we exposed 60 *D. magna* to spores derived directly from infected *Daphnia*, with 20 individuals per spore dose. For the fish-derived spores, we exposed 5 times 60 *D. magna* (20 individuals per dose) to spores derived from the faecal samples of each of five different fish. Twenty juvenile *D. magna* were not exposed to parasite spores (negative control). We individually labelled all jars and randomly distributed them across 13 trays in an incubator, rotating the tray positions weekly during medium transfers to minimise micro-environmental variation. We set the incubator to 20 °C and 16:8 h light–dark photoperiod. After a week, we topped up the medium in each jar to 90 mL. Individuals were monitored daily over four weeks for survival and offspring production to estimate castration, with visible disease symptoms emerging during the second week post-infection. We excluded from analysis individuals that died during the first week of the experiment. *Daphnia* that died after week 1 were individually crushed and examined under the microscope to check for the presence of *P. ramosa* spores.

We used the R package brglm2 (Kosmidis, 2020) to analyse binary outcomes of exposure (infected/not infected), employing generalized linear models (GLM) with a binomial distribution and bias-reduced fitting to handle potential data separation. The model included interaction terms between spore origin and spore concentration.

#### Spore production

For animals exposed to the highest spore dose, i.e. 180,000 spores, we quantified and compared the number of spores per infected host (spore load) in infected *Daphnia* treated with either *Daphnia*-derived or fish-derived spores. For this, we collected individuals on day 28 post exposure and placed them individually in 1.5-mL Eppendorf tubes with 1 mL water, stored at 4 °C. A few days later, we individually crushed the *Daphnia* using a micro-pestle and quantified the spores with a hemocytometer. We compared spore counts between treatment groups (*Daphnia*-derived vs. fish-derived spores) using Welch’s two-sample t-test to account for unequal variances between groups.

### Infection experiment from water body

We conducted infection experiments to investigate whether *D. magna* can acquire infections from *P. ramosa* spores released during feeding activity (sloppy feeding) or from spores excreted by fish not as part of the faecal droppings. Water samples with spores were collected from the previous exposure experiment: a) water that remained in the tanks for 30-40 minutes after fish feed on infected *Daphnia* (sloppy feeding), and b) water retained in the fish tanks for the two days following fish feeding.

We collected a total of 200 two-day-old *D. magna* within a 24-hour period and pooled them in a single 1.5-L jar. The next day, we retrieved the previously stored water samples from the cold room and placed them on magnetic stirrers at 750 rpm (Heidolph MR 2002) for 30 minutes. After stirring, we distributed the water samples into 200 100-mL jars, and individually transferred the juvenile *Daphnia* into these jars (each 100 jars for the water collected shortly after feeding and the water collected at the end of the two-day period) with 20 individuals assigned to each of the five fish replicates. All jars were handled and monitored as described in the previous experiment. As a negative control, we exposed 40 *Daphnia* to water from tanks where the fish were not fed infected *Daphnia*. We used one tank per treatment, with 20 *Daphnia* per group. We calculated the proportion of infected animals per treatment group and per fish replicate and used these proportions in the analysis.

We performed all statistical analyses using R Statistical Software v4.2.2 (R Core Team, 2022) within RStudio v2024.12.1.563 (RStudio Team, 2024).

## Results

### Spore recovery from gut passage

Our pilot experiment indicated that, although faeces were collected on all three days, the majority of spores were excreted within the first 24 hours post-feeding, with only occasional traces detected the following day. Afterwards, spores were no longer detectable. In the main recovery experiment, spore counts varied highly across fish individuals, but peaked within the first 24 hours post-ingestion, with most spores being excreted between 12 and 24 hours after ingestion. Spore output then dropped to low levels for the remainder of the sampling window (Fig. 1). Total recovered spore counts varied from 3.48 million to 10.5 million across individual fish, with a mean of about 7.9 million spores. This corresponded to a mean recovery rate of 60.8%, based on our estimated ingestion dose of 13 million spores per 5 infected *Daphnia* per fish.

**Figure 1.**
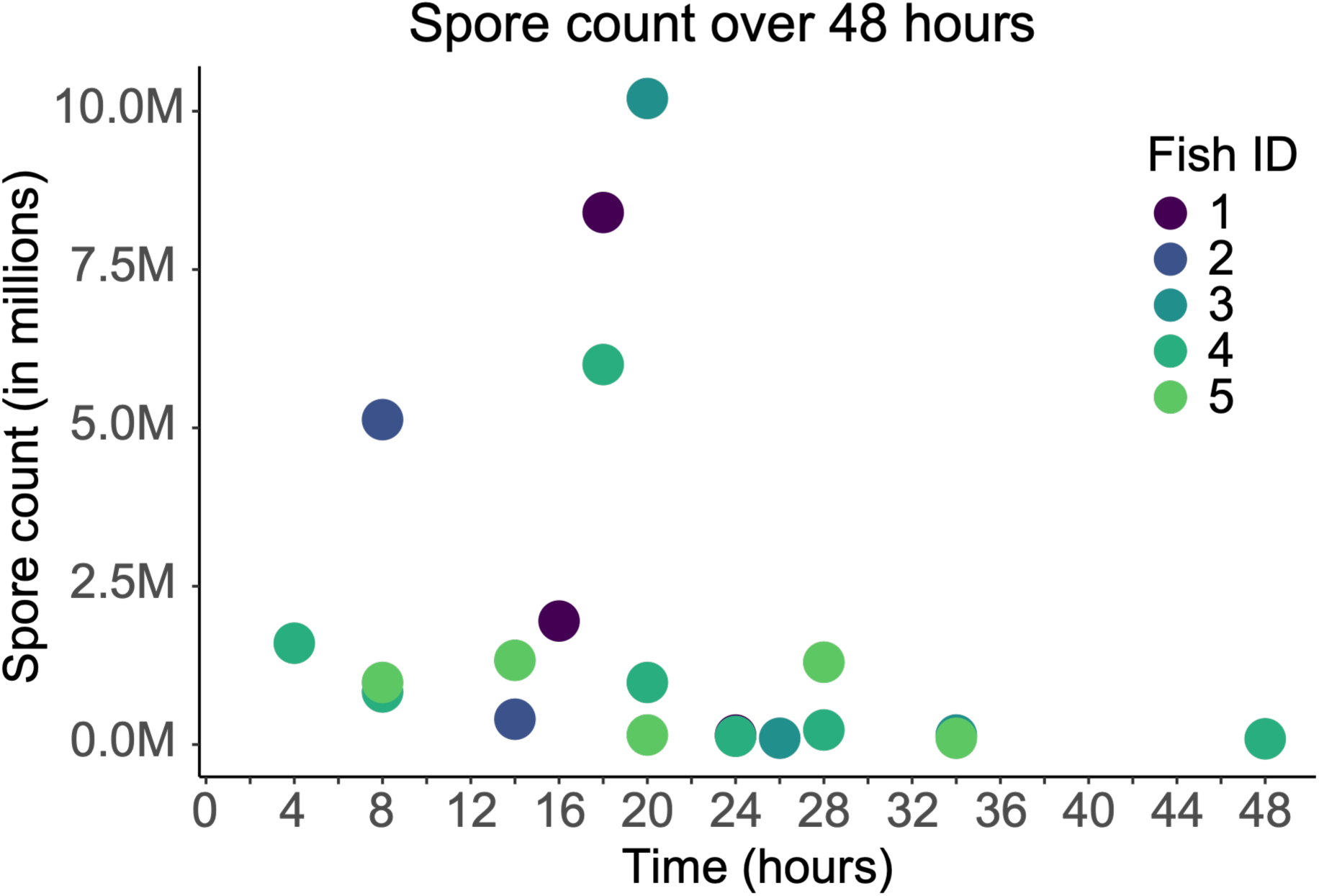
Recovery of *Pasteuria ramosa* spores from stickleback (*Gasterosteus aculeatus*) faeces over 48 hours after consumption of five infected *Daphnia magna*. Each colour represents one individual fish. Spore counts are shown in millions.

### Infection experiment with gut passaged spores

The proportion of infected hosts (infection rates) generally exceeded 70 % across all treatment combinations, often reaching 100 %. Neither spore dose nor treatment (*Daphnia*-versus fish derived spores) showed a systematic effect (Fig. 2A). All *Daphnia magna* in the negative control treatment remained uninfected throughout the experiment.

**Figure 2.**
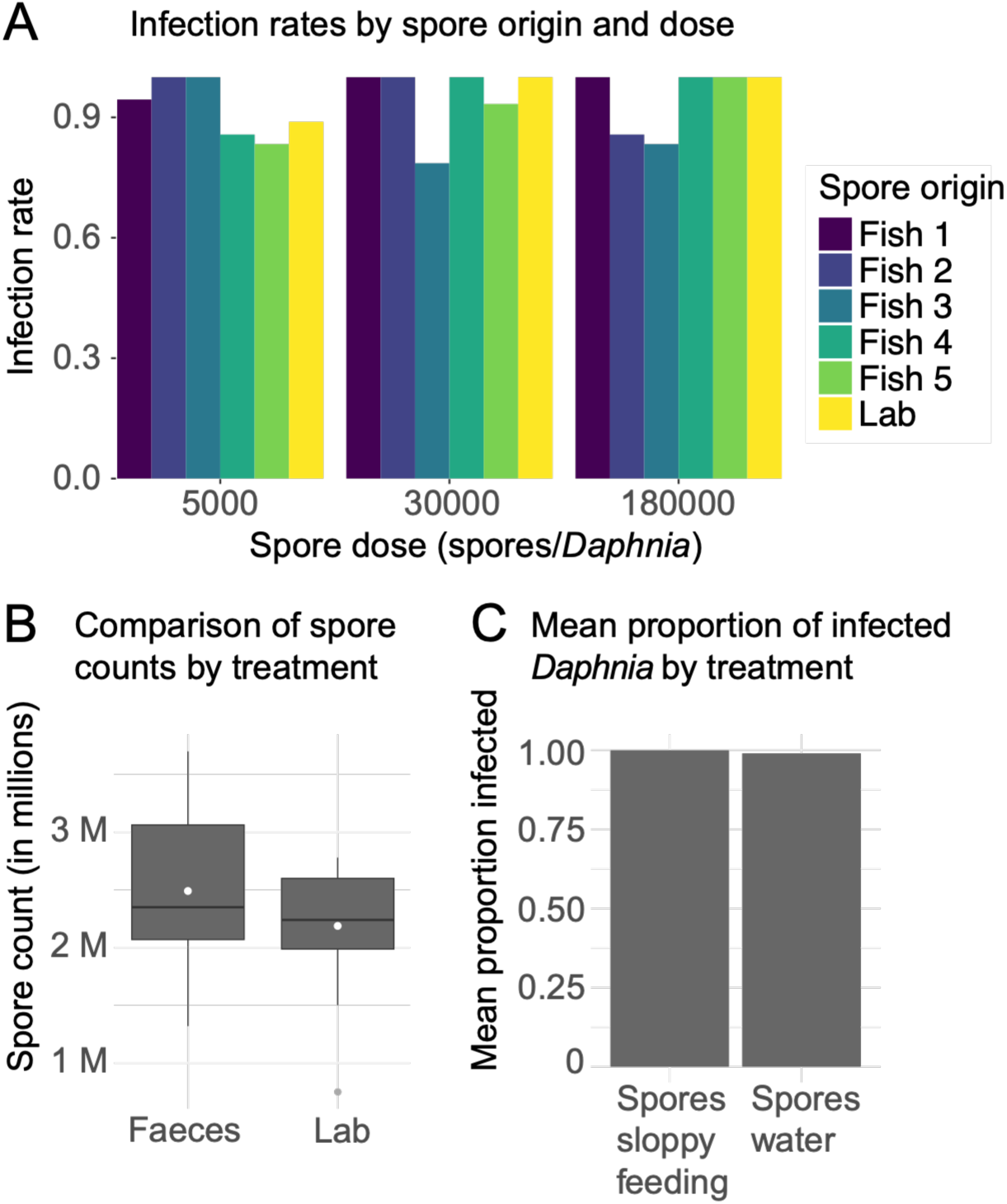
Infectivity and spore production of *Pasteuria ramosa* spores following fish-handling and gut passage. **(A)** Infection rates of *Daphnia magna* exposed to *P. ramosa* C1 spores originating either directly from infected *Daphnia* (Lab; yellow) or from stickleback faeces (Fish 1–5; light green to dark purple), across three exposure doses (5,000, 30,000, and 180,000 spores per host). Bars represent the proportion of infected individuals within each replicate group. **(B)** Parasite spore loads measured in infected *D. magna* at day 28 post-exposure to fish-derived (Faeces) or *Daphnia*-derived (Lab) spores at the highest exposure dose (180,000 spores). Boxes indicate the interquartile range (IQR); horizontal lines show medians; white points indicate means, and grey points represent values beyond 1.5 × IQR. **(C)** Infection rates of *D. magna* following exposure to water collected either shortly after fish feeding on infected *Daphnia* (spores sloppy feeding) or from tanks housing fish for two days after feeding (spores water). Bars show mean infection proportion across five replicate groups (n = 100 per treatment). Spore concentrations in water-derived treatments were not quantified.

The mean spore load of individuals exposed to high doses (180,000 spores) did not vary significantly depending on the origin (*Daphnia*-versus fish-derived spores). The faeces-derived treatment was slightly, but not significantly, higher (mean = 2,490,556) than the *Daphnia*-derived (Lab) treatment (mean = 2,188,125) (Welch’s t-test (t_37_ = 1.74, p = 0.091; Fig. 2B). All negative controls remained uninfected, with a zero spore load throughout the experiment.

### Infection experiment (spores from water vs spores from sloppy feeding)

*D. magna* exposed to *P. ramosa* spores from sloppy feeding and from post-excretion tank water exhibited uniformly high infection rates (Fig. 2C; Welch’s t-test: t(4) = 1.00, p = 0.374). Individuals in the negative control treatment showed no signs of infection.

## Discussion

Our study shows that *Pasteuria ramosa* spores retain high infectivity after passage through the gut of a predatory fish, with a high recovery rate of 61 %. Furthermore, spores released into the water during the feeding behaviour of the fish are also infective. Thus, predatory fish can be a powerful vector for *P. ramosa* parasites in freshwater ecosystems.

### Spore recovery and parasite resilience

Our spore recovery experiment confirms that a substantial proportion of *P. ramosa* spores survive gastrointestinal transit through fish and are released back into the environment via faeces. We retrieved, on average, 61% of ingested spores, most of them excreted within 24 hours after feeding. These numbers align closely with observations by Auld and Duffy (referenced in Duffy et al., 2019), who recovered approximately 67% of viable *P. ramosa* spores following gut passage through bluegill sunfish (*Lepomis macrochirus*). This rapid and efficient recovery highlights the resilience of *P. ramosa* spores, likely supported by their protective exosporium and the thick, endospore wall (Ebert et al., 1996). The high durability of these spores has been previously reported in a study by Duffy and Hunsberger (2019), who found that *P. ramosa* spores remained infective after being stored at −20°C for up to one year. Moreover, Decaestecker et al. (2004) recovered viable *P. ramosa* spores from pond sediments over 30 years old. *Pasteuria* is an endospore forming bacterium closely related to the genus *Bacillus* (Thivolle et al 2026), which has some members whose endospores survive for very long times and are used for pest controls (e.g. *B. thuringensis*) and biological warfare agents (*B. anthracis*) (Goel, 2015; Thivolle et al 2026).

The results of our study also align with the findings of Duffy (2009), who reported that approximately 50 % of spores of the fungal parasite *Metschnikowia* consumed and excreted faecally by bluegill sunfish remained infective to *Daphnia dentifera*. Although the parasite and host species differ from our system, the ecological parallel shows a broader mechanism by which non-host predators may contribute to parasite persistence.

Although *Daphnia magna* infected with *P. ramosa* have a shortened lifespan compared to uninfected hosts, they are still relatively long lived (Ebert et al., 2016). Spore counts in infected hosts increase continuously throughout their lifetime (Ebert et al., 2004; Ben-Ami et al., 2008; Hall and Ebert 2012), but transmission is only possible when the spores are released from the host’s cadaver after death. Our study shows that predation can shortcut this period and help the parasite transmit much more quickly. According to our spore recovery data, a substantial number of spores can be excreted very early in the digestive process. For example, one individual fish released nearly half of its total recovered spores (close to 6 million) within the first 8 hours (Fig. 1), showing how rapidly viable spores may be deposited back into the environment. This fast-paced excretion could accelerate transmission dynamics, exposing susceptible hosts sooner to spores and reinforcing the role of non-host predators in parasite dynamics.

According to our calculations, about a third of the ingested spores were not found in the faeces. At least a portion of these were released into the water with sloppy feeding and fish excretions; however, it is unclear if all spores survive ingestion and gut passage by fish, or if some part of them are degraded during digestion. In an experimental setup, spores may also adhere to tank walls or substrate, or to the body or mucus of the fish. Unfortunately, our design did not allow for the precise quantification of all spores in the water, limiting our ability to estimate of their fate.

### Infectivity of spores

Our study demonstrates that spores derived both directly from *Daphnia* and those recovered from fish faeces successfully infect *D. magna* at consistently high rates, even at the lowest exposure dose of 5,000 spores. The C1 *P. ramosa* isolate is known to be highly infective to the *D. magna* genotype HU-HO-2 (Hall & Ebert, 2012) and are highly resilient in maintaining their function even after exposure to digestive conditions. Moreover, no significant differences in the resulting infections were found, as the spore counts in the infected *Daphnia* did not differ significantly among treatments. These findings collectively suggest that the transmission potential of *P. ramosa* is unaffected by its route of release, whether directly from a dead host or via trophic processing in a predator.

In addition to the infection success of faeces-derived spores, we also found that spores released into the water during sloppy feeding and those present in the water post-excretion led to high infection rates in *Daphnia*. This reveals two additional and ecologically plausible transmission pathways for *P. ramosa* to reach new hosts. Given that sloppy feeding can release spores immediately during prey handling, while faecal spores may persist in the water column for extended periods, our findings underscore the environmental durability of *P. ramosa* spores and their capacity to exploit multiple routes to maintain transmission.

### Fate of spores

The identification of three transmission routes by which parasite can spread—sloppy feeding, faecal deposition, and waterborne dispersal post-defecation—expands our understanding of parasite epidemiology within freshwater ecosystems. These pathways represent distinct, yet complementary mechanisms for reinfecting susceptible hosts, allowing *P. ramosa* to exploit delayed and spatially disconnected transmission opportunities.

From an ecological standpoint, these dynamics imply that predator–prey interactions can simultaneously suppress host density *and* promote parasite spread, highlighting a more complex role for predators in disease ecology: Rather than functioning solely as parasite sinks, predators such as *G. aculeatus* may also act as parasite vectors, depending on ecological context.

Faecal deposition of spores in littoral zones, areas characterized by high biological activity and shared by foraging fish and *Daphnia* (Schindler & Scheuerell, 2002; Vadeboncoeur et al., 2011; Arbore et al., 2016), creates important infection interfaces. Spores deposited in these zones are likely to be ingested by *Daphnia* feeding on algae and detritus (Rautio & Vincent, 2006). These sediments can act as long-term reservoirs for viable spores, which may later be resuspended into the water column through abiotic forces such as wave action and wind (Li et al., 2017), or by biotic activity (Scheffer et al., 2003). This resuspension enhances spore availability and can increase transmission opportunities when overlapping with *Daphnia* habitats.

Several studies have also recently reported that *P. ramosa* spores can be recovered from the free water (Shaw and Duffy, 2023; Shaw et al., 2024; Tadiri and Ebert, 2026), a surprising result, as spores were believed to be released only from dead hosts, which quickly sink to the bottom of the water body and result in sediment borne transmission stages (Tadiri & Ebert 2026). Here we show that, in addition to sediment-associated dispersal, sticklebacks can spread viable spores directly into the open water through sloppy feeding and liquid excretion, enabling pelagic dispersal. Tadiri and Ebert (2026) estimates that spores in the free water sink only very slowly to the bottom, allowing for a considerable time period of exposure of pelagic parasite spores to filter feeding *Daphnia*. These processes increase the distribution of infectious stages beyond the benthic zone, enhancing the parasite’s ability to exploit multiple spatial niches within freshwater ecosystems.

Beyond local transmission, fish can contribute to broader parasite dispersal. *Daphnia* inhabit lentic water bodies, whereas fish such as sticklebacks may also occupy lotic environments, including streams and rivers. In lake systems connected by these flowing waters, fish movements can transport *P. ramosa* spores from one lake to another, even against the direction of water flow, thereby contributing to the long-distance spread of the parasite across freshwater networks.

Ultimately, the transmission landscape of *P. ramosa* is influenced by the extent of spatial and temporal overlap between fish faecal deposition sites, spore dispersal mechanisms, and *Daphnia* microhabitats shaped by environmental factors and species behaviours. This highlights the importance of considering not only direct predator-prey interactions but also indirect dispersal pathways that govern how infective spores reach and infect susceptible hosts across freshwater ecosystems.

## Conclusion

This study demonstrates that fish can act as vectors for the bacterial parasite *Pasteuria ramosa*, influencing its transmission dynamics to *Daphnia magna* by adding additional routes of transmission to the previous picture. This contradicts a core assumption of the healthy herd hypothesis, namely that predation eliminates parasites from the population and therefore reduces the potential of infectious diseases to spread. Thus, our findings reveal a more complex role for predators in aquatic disease dynamics, where fish may contribute to the spatial redistribution and environmental persistence of infectious stages in freshwater ecosystems (compare Lopez et al., 2023). Understanding the spatial and temporal overlap of predators, hosts, and infective stages under realistic ecological conditions will be essential for predicting infection risk and modelling disease epidemiology in natural freshwater ecosystems.

Our research reframes the role of predators from simple consumers to ecological participants in disease transmission. By revealing the varied routes through which parasites persist and spread in the presence of predators, this study contributes to a deeper understanding of host–parasite–predator interactions and highlights the need for integrative approaches to freshwater disease ecology.

## Acknowledgments

We thank Michelle Krebs, Jürgen Hottinger, and Urs Stiefel for their technical support. We are grateful to all members of the Ebert Research Group for creating a collaborative and intellectually stimulating environment. We also want to thank the members of the Salzburger Group, including Daniel Berner, for giving us the opportunity to work with the sticklebacks in their aquarium facilities. We thank Adrian Indermaur and Attila Rüegg for technical support and help with the fish husbandry and the aquaria. We want to thank Meghan Duffy for critical comments on the manuscript. We thank Suzanne Zweizig for help with the language. This work was supported by the Swiss National Science Foundation grant numbers 310030_188887 and 310030_219529 to D.E.

## Data availability statement

All data and code supporting the findings of this study are publicly available in the GitHub repository archived on Zenodo (Lampadaridis et al., 2026, DOI: 10.5281/zenodo.24403292). The repository contains the R scripts and associated data files used for the analysis of infection experiments, spore production, spore-crushing treatments, and related biological measurements.

## Notes

https://github.com/Carlos-Manuel-Herrera-Castillo/Planktivorous_Fish_As_Vectors

### Competing Interest Statement

The authors have declared no competing interest.

### Summary of Updates

The manuscript has been updated, including language review, updating further references, including a data availability statement and updating the acknowledgments section.

