## Supplementary Fig. 1 for "Assessing planktivorous fish as vectors of a plankton parasite"

### Supplementary material

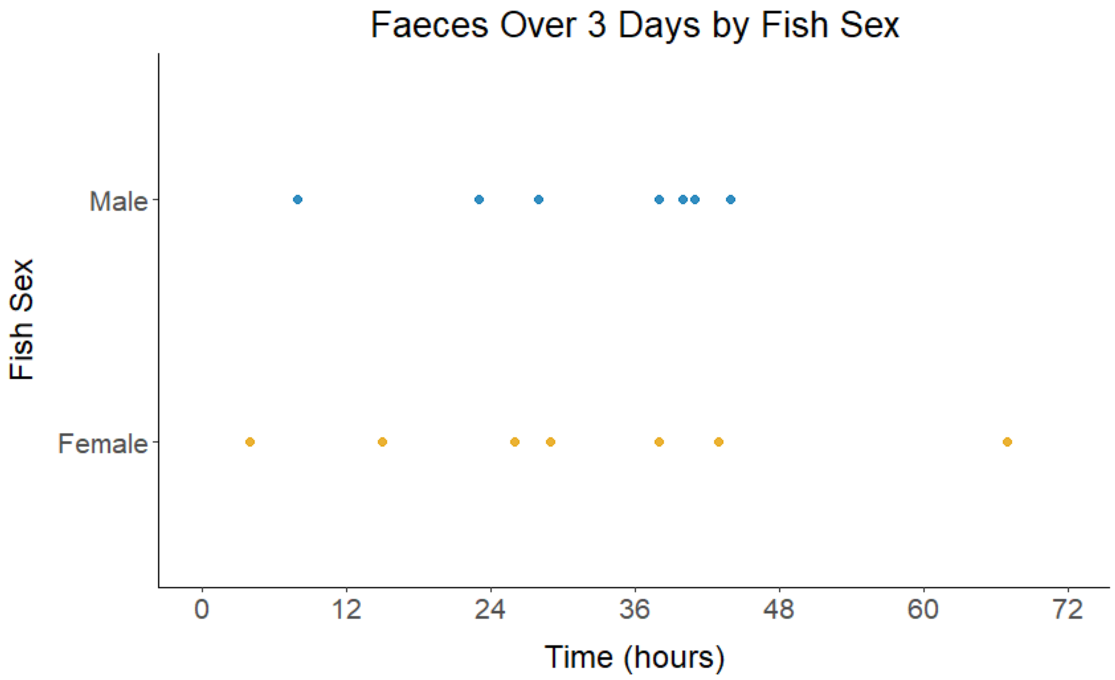


**Supplementary figure 1.** Timing of faecal deposition by a male (blue) and a female (orange) three-spined sticklebacks over a period of 3-days post feeding. Each point represents an observed faecal event, recorded at a specific time post-feeding.
